# Rapid shift in substrate utilization driven by hypothalamic Agrp neurons

**DOI:** 10.1101/086348

**Authors:** João Paulo Cavalcanti-de-Albuquerque, Marcelo R. Zimmer, Jeremy Bober, Marcelo O. Dietrich

## Abstract

Agrp neurons drive feeding. To what extend these neurons participate in the regulation of other homeostatic processes is not well understood. We investigated the role of Agrp neurons in substrate utilization in mice. Activation of Agrp neurons was sufficient to rapidly increase RER and carbohydrate utilization, while decreasing fat utilization. These metabolic changes were linearly correlated with carbohydrates ingested, but not protein or fat ingestion. However, even in the absence of ingestive behaviors, activation of Agrp neurons led to changes in substrate utilization in well-fed mice. These effects were coupled to metabolic shifts towards lipogenesis. Inhibition of fatty acid synthetase (FAS) blunted the effects of Agrp neurons on substrate utilization. Finally, Agrp neurons controlled peripheral metabolism, but not food intake, via ß3-adrenergic receptor signaling in fat tissues. These results reveal a novel component of Agrp neuron-mediate metabolism regulation that involves sympathetic activity on fat compartments to shift metabolism towards lipogenesis.

## Introduction

Obesity is a major health problem that results from altered regulation of energy metabolism. The central nervous system tightly controls energy metabolism by regulating hormonal and autonomic action on peripheral tissue. The hypothalamus is a conserved region in the brain involved in homeostatic control, including energy balance. The arcuate nucleus of the hypothalamus contains a population of neurons that selectively expresses agouti-related protein (AgRP; hereafter, Agrp neurons) (Broberger et al., 1998; Hahn et al., 1998; Ollmann et al., 1997; Rossi et al., 1998). Agrp neurons are located in proximity to the third ventricle and have direct access to circulation (Olofsson et al., 2013), as this brain region lacks a complete blood-brain barrier (Broadwell et al., 1983; Broadwell and Brightman, 1976). Consistently, Agrp neurons are known to respond to a variety of circulating factors (Könner et al., 2007; Pinto et al., 2004; Steculorum et al., 2015; van den Top et al., 2004). However, recent evidence demonstrates that Agrp neurons also integrate food-related information via sensory pathways (Betley et al., 2015; Chen et al., 2015; Mandelblat-Cerf et al., 2015). Thus, Agrp neurons are unique as they gather relevant information from several sources to regulate physiology and behavior.

Agrp neurons were first described to be active during food deprivation, a phenomenon conserved across rodent and non-human primate species (Grove et al., 2003; Hahn et al., 1998; Mandelblat-Cerf et al., 2015; Takahashi and Cone, 2005). Because Agrp neurons co-express NPY (Broberger et al., 1998; Hahn et al., 1998), and because NPY acts as a potent orexigenic peptide when injected into the brain (Clark et al., 1984), it was natural to conclude that during food deprivation Agrp neurons are active to drive food intake. Subsequent work demonstrated that activation of Agrp neurons is sufficient to drive food intake in sated mice (Aponte et al., 2011; Dietrich et al., 2015; Krashes et al., 2011). Conversely, elimination of Agrp neurons in the adult brain led to aphagia (Gropp et al., 2005; Luquet et al., 2005).

Recent work demonstrated a specific role for Agrp neurons in metabolic processes, including the control of white adipose tissue (WAT) browning and thermogenesis (Ruan et al., 2014), as well as brown adipose tissue (BAT) glucose metabolism (Steculorum et al., 2016). Here, we investigate the role of Agrp neurons in the control of peripheral substrate utilization. Our data demonstrate that Agrp neuron activation rapidly regulates peripheral metabolism independently of food ingestion. Specifically, Agrp neuron activation promotes lipogenesis via sympathetic nervous system (SNS). These results highlight the complex function of these key hypothalamic neurons in normal physiology and in disordered metabolic states, such as obesity.

## Results

### Acute switch in nutrient utilization upon Agrp activation

Nutrient utilization can be measured by indirect calorimetry, where the measurements of VCO_2_ production and VO_2_ consumption are used to calculate the respiratory exchange ratio (RER) (Figure 1A) (Frayn, 1983). Under normal conditions, a RER approaching 0.7 indicates predominant fat oxidation, while a RER approaching 1.0 indicates predominant carbohydrate oxidation (Figure 1A) (Frayn, 1983). To test for the acute role of Agrp neurons in nutrient utilization we took advantage of an animal model that we have recently characterized (Dietrich et al., 2015; Ruan et al., 2014), in which Agrp neurons are transiently activated by peripheral injection of capsaicin in *Agrp*^Trpv1^ mice (Dietrich et al., 2015; Ruan et al., 2014). *Agrp*^Trpv1^ mice are generated by crossing *Agrp*^Cre^ to *Rosa26*^LSL-Trpv1^ mice backcrossed to a *Trpv1*^KO^ background to prevent capsaicin action on other cell types (Arenkiel et al., 2008; Guler et al., 2012). As a result, *Agrp*^Trpv1^ mice selectively express the capsaicin-sensitive channel, Trpv1, in Agrp neurons. Because capsaicin is a highly specific ligand of Trpv1 (Caterina et al., 1997), peripheral injection of capsaicin allows for a rapid, reliable and transient chemogenetic activation of Agrp neurons in *Agrp*^Trpv1^ mice (Dietrich et al., 2015; Ruan et al., 2014). Using *Agrp*^Trpv1^ mice allowed us to rapidly activate Agrp neurons without the necessity of tethers (as for example, using optogenetics), facilitating the study of animals in indirect calorimetry chambers to measure gas (O_2_ and CO_2_) exchange. Importantly, Trpv1-mediated activation of Agrp neurons is both rapid (latency to start is ~2 min) and short lived (lasts ~1h), unlike the effects of activating Agrp neurons via the designer receptor hM3Dq (Krashes et al., 2011).

**Figure 1:**
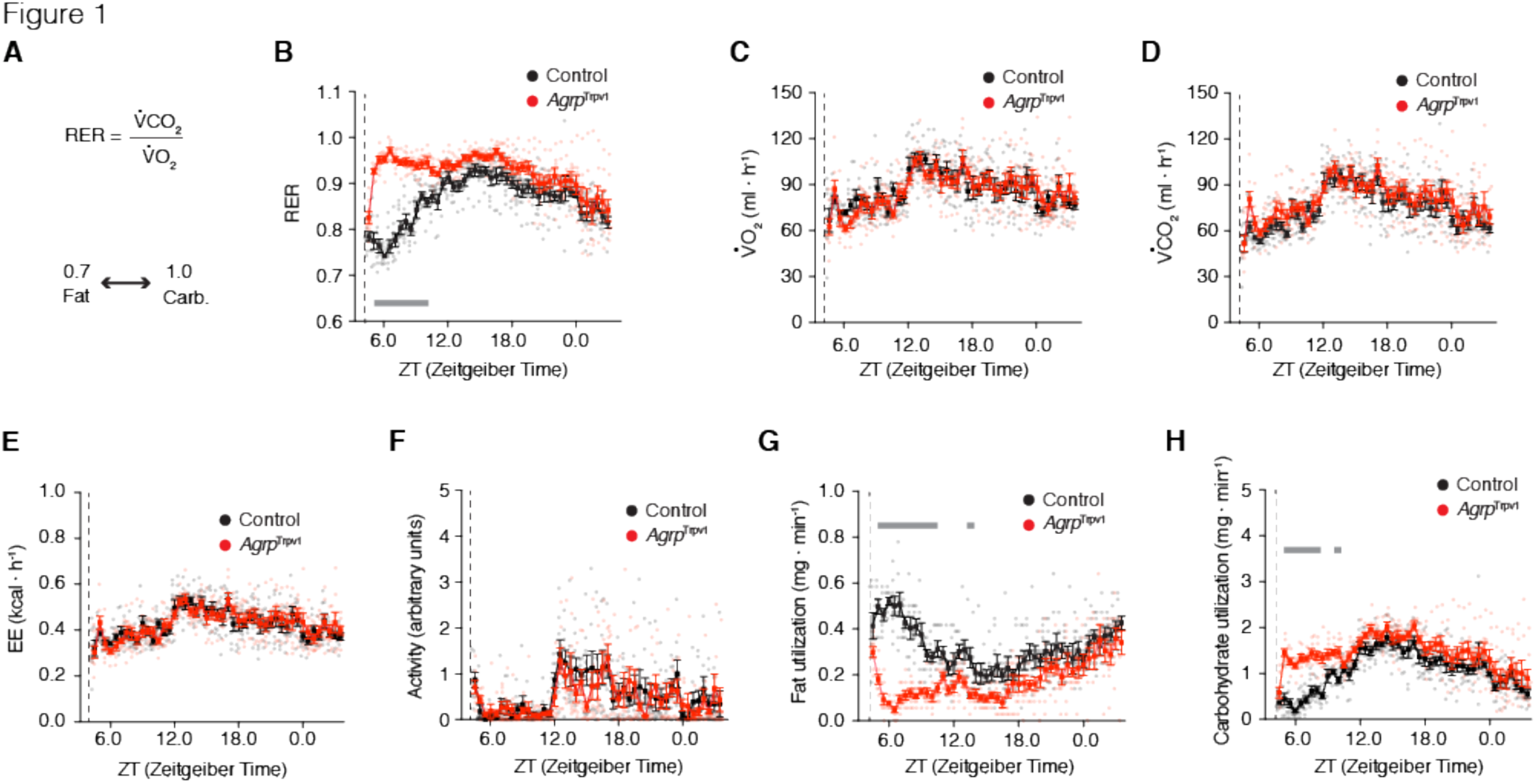
Rapid shift in substrate utilization upon activation of Agrp neurons. (**A**) In indirect calorimetry chambers, RER was calculated by diving the VCO_2_ by the VO_2_; oxidation of fat acids (e.g., palmitate) generates a RER of 0.7, while oxidation of carbohydrates (e.g., glucose) generates a RER of 1.0. From **B-H**, control (black; n = 8) and *Agrp*^Trpv1^ mice (red; n = 8) were acclimated to calorimetry chambers before been injected with capsaicin (dashed lines; 10 mg/kg, i.p.) during the light cycle with food and water provided *ad libitum*. (**B**) RER (interaction: *F*_45, 630_ = 8.40, *P* < 0.0001; time: *F*_45, 630_ = 13.86, *P* < 0.0001; group: *F*_1, 14_ = 52.59, *P* < 0.0001). (**C**) VO_2_ (interaction: *F*_45, 630_ = 1.07, *P* = 0.34; time: *F*_45, 630_ = 11.22, *P* < 0.0001; group: *F*_1, 14_ = 0.01, *P* = 0.89). (**D**) VCO_2_ (interaction: *F*_45, 630_ = 0.88, *P* = 0.68; time: *F*_45, 630_ = 15.07, *P* < 0.0001; group: *F*_1, 14_ = 1.58, *P* = 0.22). (**E**) Energy expenditure (interaction: *F*_45, 630_ = 0.96, *P* = 0.53; time: *F*_45, 630_ = 12.09, *P* < 0.0001; group: *F*_1, 14_ = 0.02, *P* = 0.87). (**F**) Ambulatory activity (interaction: *F*_45, 630_ = 0.80, *P* = 0.81; time: *F*_45, 630_ = 6.02, *P* < 0.0001; group: *F*_1, 14_ = 0.31, *P* = 0.58). (**G**) Calculated fat utilization (interaction: *F*_45, 630_ = 6.44, *P* < 0.001; time: *F*_45, 630_ = 7.22, *P* < 0.0001; group: *F*_1, 14_ = 33.83, *P* < 0.0001). (**H**) Calculated carbohydrate utilization (interaction: *F*_45, 630_ = 2.77, *P* < 0.0001; time: *F*_45, 630_ = 18.48, *P* < 0.0001; group: *F*_1, 14_ = 16.73, *P* = 0.001). Statistical analysis was performed using two-way ANOVA with time as a repeated measure followed by Holm-Sidak’s multiple comparisons test (MCT). Grey bars indicate time points in which MCTs were statistically significant (*P* < 0.05). Dashed line indicates time of capsaicin injection. Small symbols indicate individual values. Large symbols indicate mean ± SEM.

We injected mice with capsaicin during the light cycle and recorded concomitant changes in locomotor activity and gas exchange for the ensuing 24 hours. Capsaicin injections produced sharp increases in RER in *Agrp*^Trpv1^ mice, (Figure 1B), in line with the previously observed fast kinetics of feeding induced by Agrp neuronal activation in this animal model (Dietrich et al., 2015). We did not observe statistically significant changes in either VO_2_ (Figure 1C), VCO_2_ (Figure 1D) or energy expenditure (Figure 1E). Because exercise increases RER, we also measured activity levels upon Agrp neuron activation. In line with our previous report (Dietrich et al., 2015), activation of Agrp neurons in the home cage in the presence of food did not increase levels of ambulatory activity (Figure 1F), indicating that the increase in RER was not affected by elevated locomotion. Based on gaseous exchange (Frayn, 1983), we calculated total rates of fat utilization and carbohydrate utilization for the whole animal. In line with changes in RER, activation of Agrp neurons led to a rapid and prolonged decrease in fat utilization (Figure 1G) concomitant with an increase in carbohydrate utilization (Figure 1H). Because (i) this experiment was performed in the presence of food and (ii) the effects on nutrient utilization were prolonged compared to feeding upon Agrp neuron activation (Dietrich et al., 2015), we hypothesized that these metabolic shifts are due to changes in postprandial metabolism.

### Effects of diet ingestion on metabolic switches

To further dissect the effects of altered RER from increased food intake upon Agrp neuron activation, we determined how diet composition affects the metabolic phenotypes by varying the proportion of macronutrients in the diet (Figure 2A; see Material and Methods for more details). We activated Agrp neurons in *Agrp*^Trpv1^ mice during the light cycle, providing controlled amounts of different diets: LFHS, a low-fat high-sugar diet containing 10% kcal from fat sources; HF45, a high-fat diet containing 45% kcal from fat sources; and HF60, a high-fat diet containing 60% kcal from fat sources. The three diets contain 20% kcal from protein sources. We performed these experiments in three different conditions: (i) limiting the amount of calories ingested from fat sources (0.4 kcal from fat, ‘Fat Clamp’); (ii) limiting the amount of calories ingested from carbohydrate sources (0.7 kcal from sugars, ‘Sugar Clamp’), or (iii) feeding animals isocaloric quantities of food (1.5 kcal from all sources, ‘Cal Clamp’).

**Figure 2:**
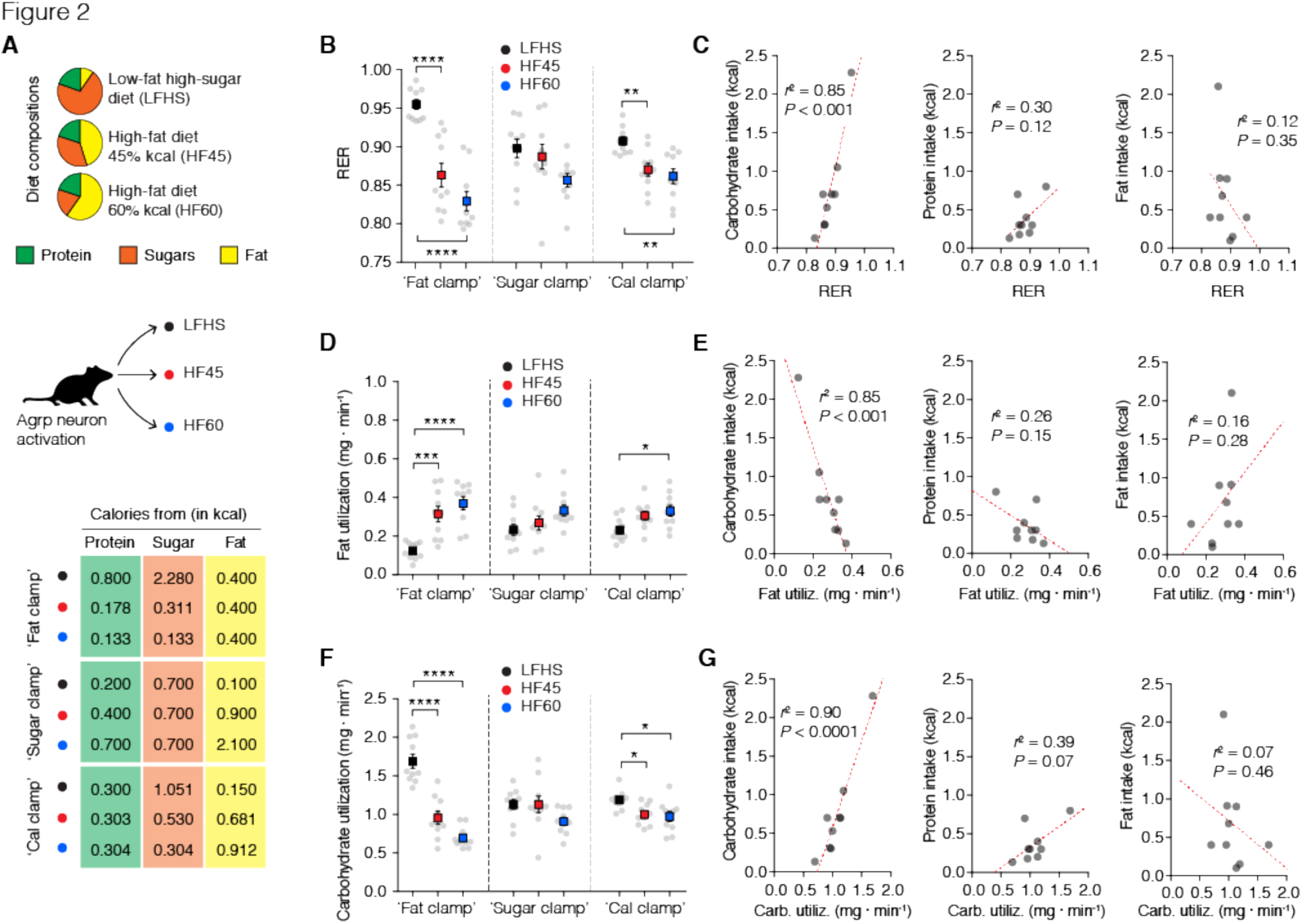
Substrate utilization in response to different diets. Mice (*Agrp*^Trpv1^) were tested in calorimetry chambers for their rapid response to different diets upon activation of Agrp neurons (with capsaicin, 10 mg/kg, i.p). (**A**) Three different diets were used in the studies, containing different distribution of macronutrients from similar sources (protein levels were equal at 20 kcal%); low-fat high-sugar diet (LFHS; Research Diets D12450B [carbohydrates: 70 kcal%; fat: 10 kcal%]); high-fat diet (HF45; Research Diets D12451 [carbohydrates: 35 kcal%; fat: 45 kcal%]); and high-fat diet (HF60; Research Diets D12492 [carbohydrates: 20 kcal%; fat: 60 kcal%]). Mice were fed pre-weighted amounts of the three different diets in three different experimental conditions (see table; n = 10 mice per condition): ‘Fat clamp’, all mice received a pellet of the diet containing 0.4 kcal from fat; ‘Sugar clamp’, mice received a food pellet containing 0.7 kcal from carbohydrates; and ‘Cal clamp’, mice received a food pellet containing a total of 1.5 kcal from all sources. (**B**) Mean RER after activation of Agrp neurons (related to Figure 2 supplement 1A): ‘Fat clamp’ (*F*_2, 27_ = 29.13, *P* < 0.0001); ‘Sugar clamp’ (*F*_2, 27_ = 2.662, *P* = 0.08), and ‘Cal clamp’ (*F*_2, 27_ = 4.753, *P* = 0.01). (**C**) Correlation between RER (abscissa) and macronutrient intake (ordinate). (**D**) Mean calculated fat utilization after activation of Agrp neurons (related to Figure 2 supplement 1F): ‘Fat clamp’ (*F*_2, 27_ = 16.5, *P* < 0.0001); ‘Sugar clamp’ (*F*_2, 27_ = 2.878, *P* = 0.07), and ‘Cal clamp’ (*F*_2, 27_ = 8.897, *P* = 0.001). (**E**) Correlation between fat utilization (abscissa) and macronutrient intake (ordinate). (**F**) Mean calculated carbohydrate utilization after activation of Agrp neurons (related to Figure 2 supplement 1G): ‘Fat clamp’ (*F*_2, 27_ = 45.83, *P* < 0.0001); ‘Sugar clamp’ (*F*_2, 27_ = 2.627, *P* = 0.09), and ‘Cal clamp’ (*F*_2, 27_ = 5.53, *P* = 0.009). (**G**) Correlation between carbohydrate utilization (abscissa) and macronutrient intake (ordinate). In B, D and F, ordinary one-way ANOVA was used to test for statistical differences within each experimental condition (‘Clamp’) followed by Holm-Sidak’s multiple comparisons test (MCT). MCTs are indicated as * *P* < 0.05, ** *P* < 0.01, *** *P* < 0.001, and **** *P* < 0.0001. Large symbols indicate mean ± SEM. Small grey symbols indicate individual values. Ten mice were used per condition. In C, E, and G, linear regression analysis was used to calculate the correlation between total macronutrient intake and substrate utilization measurements (as plotted in B, D, and F); each point represents the mean response of 10 animals in the same condition (from B, D, and F); *r*^2^ and *P* values are plotted in each panel; dashed red line represents the linear regression model. Figure 2 supplement 1 presents detailed analysis of the calorimetry data and activity levels.

Before activation of Agrp neurons, baseline RER was 0.79 ± 0.003 (mean ± s.e.m.; n = 90 measurements), fat utilization was 0.51 ± 0.01 mg • min^−1^ (mean ± s.e.m.; n = 90 measurements) and carbohydrate utilization was 0.61 ± 0.02 mg • min^-1^ (mean ± s.e.m.; n = 90 measurements), indicating that in the conditions tested animals were using substrates from mixed sources. Following activation of Agrp neurons, we observed an increase in RER in all conditions tested (Figure 2B and Figure 2 - Figure supplement 1A). In linear regression analyses, we found a strong correlation between RER and carbohydrate ingestion (r^2^ = 0.85, *P* < 0.001), but not protein (r^2^ = 0.30, *P* = 0.12) or fat (r^2^ = 0.12, *P* = 0.35) (Figure 2C). Fat utilization was reduced upon Agrp neuron activation in all conditions tested (Figure 2D and Figure 2 – Figure supplement 1F) and was negatively correlated with carbohydrate ingestion (r^2^ = 0.85, *P* < 0.001), but not protein (r^2^ = 0.26, *P* = 0.15) or fat (r^2^ = 0.16, *P* = 0.25) (Figure 2E). Carbohydrate utilization followed changes in RER, increasing upon Agrp neuron activation in all dietary conditions tested (Figure 2F and Figure 2 - Figure supplement 1G). Changes in carbohydrate utilization were strongly correlated to carbohydrate ingestion (r^2^ = 0.90, *P* < 0.0001; Figure 2G). Protein ingestion also correlated with carbohydrate utilization, but linear regression analysis did not reach statistical significance (r^2^ = 0.39, *P* = 0.07; Figure 2G). Fat intake did not correlate with carbohydrate utilization (r^2^ = 0.07, *P* = 0.46; Figure 2G). As expected, these results demonstrated that carbohydrate levels in the diet are main players in the metabolic shift that occurs upon food ingestion, suggesting that macronutrient quality rather than caloric content account for these effects.

### Agrp neurons control substrate utilization in the absence of ingestion

The effect of Agrp neuron activation on carbohydrate metabolism led us to further explore the extent to which activation of Agrp neurons and sugar ingestion interact to rapidly shift metabolism. We provided different concentrations of glucose to mice via gavage delivery, thereby ruling out potential cephalic phase effects (Figure 3A). Gavage infusion of glucose led to dose-dependent increases in RER (Figure 3B; r^2^ = 0.99, *P* = 0.003), decreases in fat utilization (Figure 3C; r^2^ = 0.99, *P* = 0.004), and increases in carbohydrate utilization (Figure 3D; r^2^ = 0.99, *P* = 0.003). These metabolic shifts were mainly due to increases in VCO_2_ (Figure 3F; r^2^ = 0.98, *P* = 0.006) and not VO_2_ (Figure 3E; r^2^ = 0.84, *P* = 0.07). Glucose intake also positively correlated with energy expenditure (Figure 3G; r^2^ = 0.92, *P* = 0.03), likely due to the thermic effects of carbohydrate digestion. Activity levels were unchanged (Figure 3D; r^2^ = 0.58, *P* = 0.23). These experiments demonstrate that small amounts of carbohydrate ingestion alone are sufficient to shift metabolism even in the absence of carbohydrate sensing at the level of the mouth. Thus, our previous results on substrate metabolism upon activation of Agrp neurons could be simply explained by ingestion of carbohydrates. We designed the next experiments to test this consideration.

**Figure 3:**
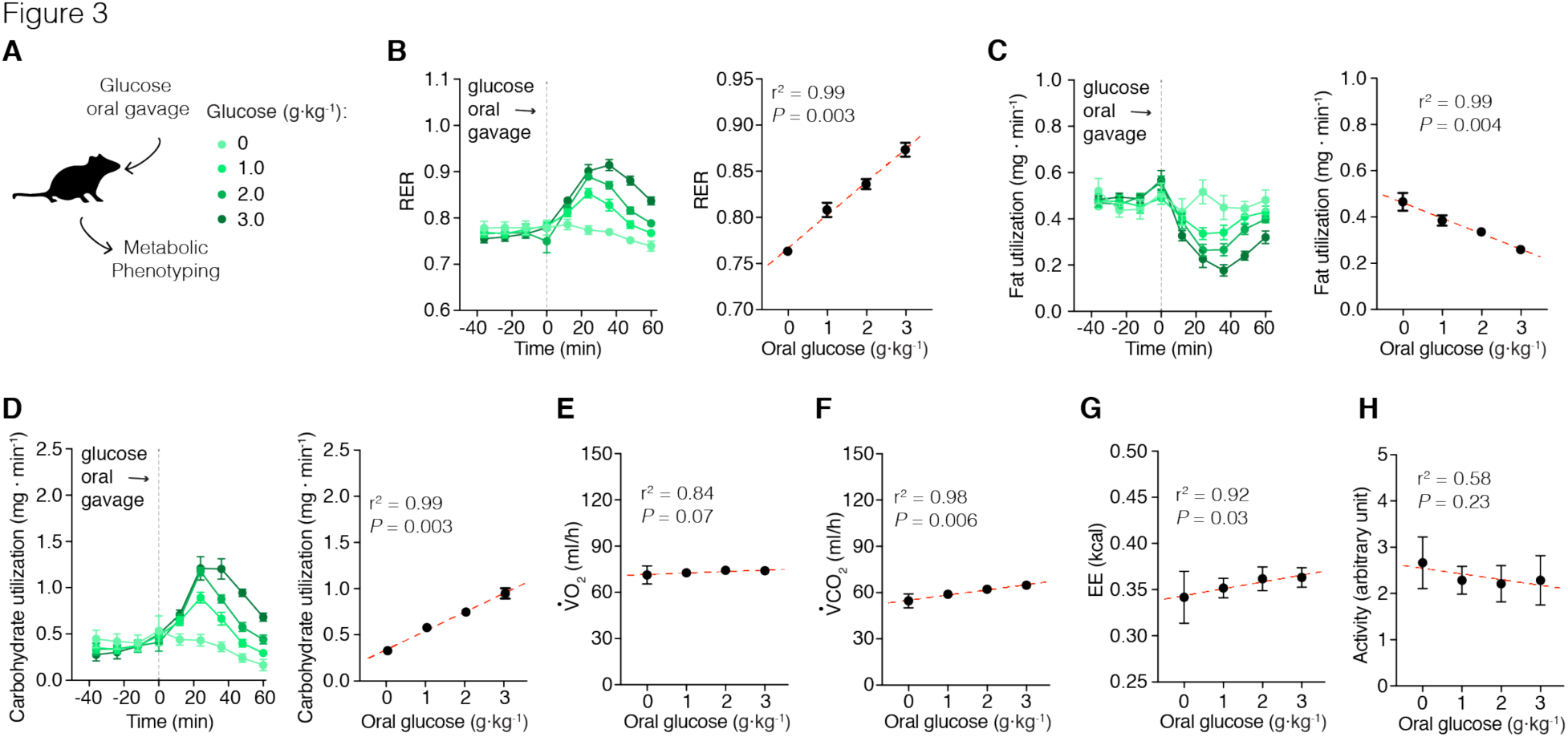
Acute effects of glucose on metabolism. Metabolic phenotyping of mice immediately after a dose-response of glucose (via gavage). (**A**) Mice received a bolus of saline or glucose solution (1, 2, or 3 g/kg body weight dissolved in saline) via gavage. (**B**) Glucose gavage infusion produces an acute increase in RER that is dose dependent. (**C**) Calculated fat utilization negatively correlates to glucose ingestion. (**D**) Calculated carbohydrate utilization is positively correlated to glucose ingestion. (**E**) VO_2_ levels did not correlated with glucose infusion. (**F**) VCO_2_ was positively correlated with glucose ingestion. (**G**) Glucose gavage also positively correlated with energy expenditure measurements, but did not correlated with changes in (**H**) activity levels. The total number of animals used in this study was: saline (n = 7); glucose 1 g/kg (n = 12); glucose 2 g/kg (n = 16); and glucose 3 g/kg (n = 8). Symbols represent mean ± SEM. Grey dashed line indicates time of oral gavage. In the linear correlation panels, symbols indicate mean of all mice in the given group ± SEM; dashed red line represents the linear regression model;*r*^2^ and *P* values are plotted in each panel.

We infused both control and *Agrp*^Trpv1^ mice with glucose (2 g/kg body weight, via gavage) and activated Agrp neurons by injecting capsaicin (Figure 4A). Strikingly, activation of Agrp neurons led to an increased peak and a more sustained elevation of RER (Figure 4B), an effect that was present even in mice infused with saline (Figure 4B). Accordingly, activation of Agrp neurons led to a sustained decrease in fat utilization (Figure 4C) and increase in carbohydrate utilization (Figure 4D), regardless of glucose infusion. These results strongly suggest that activation of Agrp neurons alone is sufficient to promote shifts in substrate utilization, independently of food consumption. However, even in animals infused with saline, small amounts of liquid were delivered via gavage, raising the possibility that gastric distension acts together with Agrp neuron activation to promote changes in substrate utilization. To exclude this possibility, we repeated our experiments in a new cohort of mice in which Agrp neurons were activated in the absence of food (Figure 4 supplement 1A). In line with our previous findings, activation of Agrp neurons alone was sufficient to increase RER (Figure 4 supplement 1B), while decreasing fat utilization (Figure 4 supplement 1G) and increasing carbohydrate utilization (Figure 4 supplement 1H). The above experiments demonstrate that Agrp neurons rapidly control whole body substrate utilization by shifting metabolism towards carbohydrate relative to fat utilization independently of ingestive behaviors.

**Figure 4:**
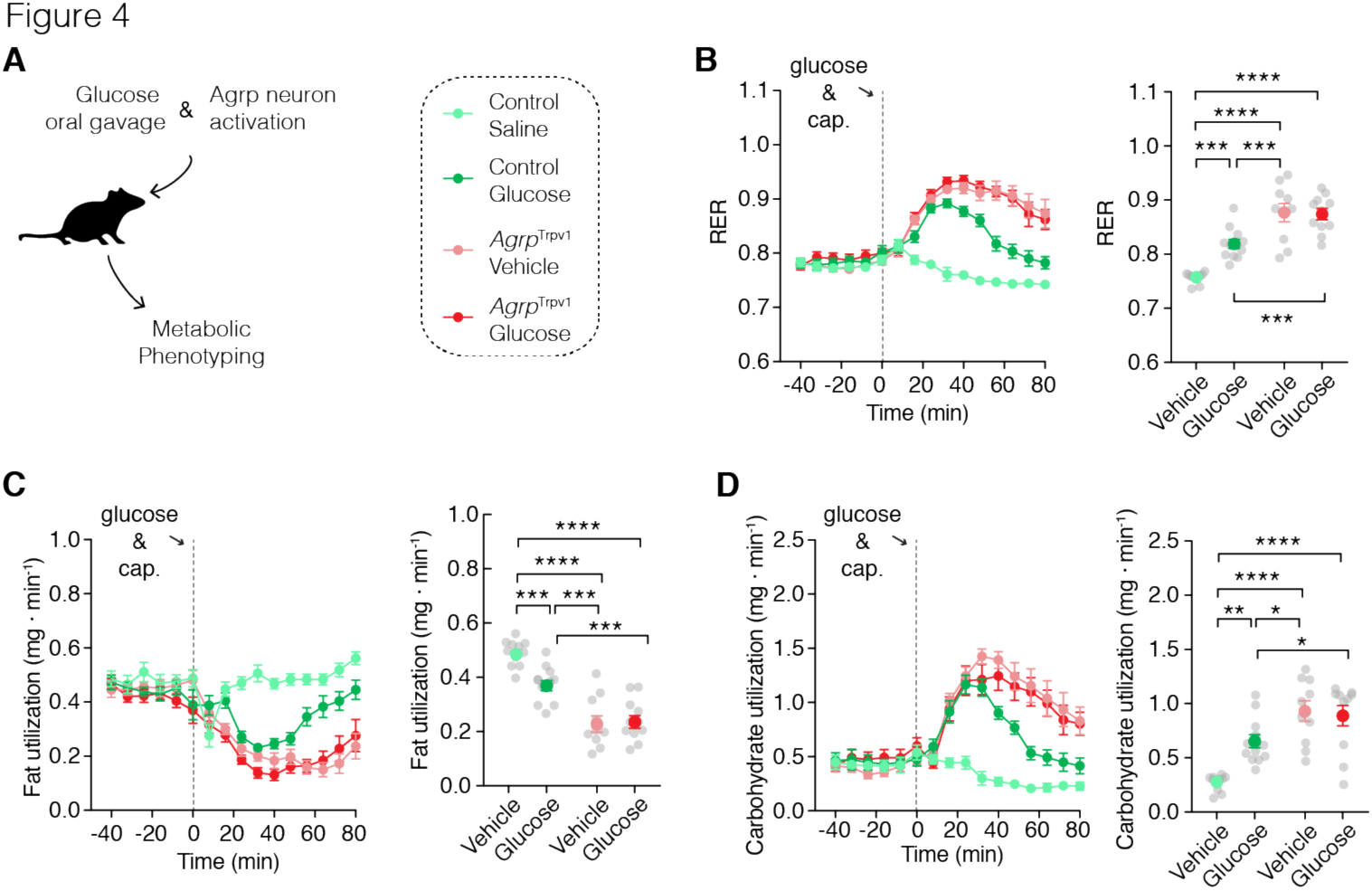
Agrp neurons control substrate utilization independently of ingestion. (**A**) Control and *Agrp*^Trpv1^ mice received a bolus of saline or glucose (2 g/kg) via gavage followed by peripheral injection of capsaicin (10 mg/kg, i.p.). (**B**) Changes in RER in control mice fed different doses of glucose solutions (interaction: *F*_1, 40_ = 9.82, *P* = 0.003; gavage solution: *F*_1, 40_ = 8.20, *P* = 0.006; genotype: *F*_1, 40_ = 71.31, *P* < 0.0001). (**C**) Fat utilization (interaction: *F*_1, 40_ = 7.91, *P* = 0.007; gavage solution: *F*_1, 40_ = 6.07, *P* = 0.01; genotype: *F*_1, 40_ = 79.82, *P* < 0.0001). (**D**) Carbohydrate utilization: (interaction: *F*_1, 40_ = 8.29, *P* = 0.006; gavage solution: *F*_1, 40_ = 5.26, *P* = 0.02; genotype: *F*_1, 40_ = 37.44, *P* < 0.0001). Statistical analysis was performed using two-way ANOVA on the mean response after gavage and capsaicin injection; genotype (control vs. *Agrp*^Trpv1^) and gavage infusion (saline vs. glucose) were used as factors for the ANOVA. Holm-Sidak’s multiple comparisons test (MCT) was used to find post-hoc differences among groups. MCTs are indicated as * *P* < 0.05, ** *P* < 0.01, *** *P* < 0.001, and **** *P* < 0.0001 in figure panels. Control mice + saline gavage (n = 11); control mice + glucose gavage (n = 12); *Agrp*^Trpv1^ mice+ saline gavage (n = 10); *Agrp*^Trpv1^ mice + glucose gavage (n = 11). Dashed grey line indicates time of oral gavage and capsaicin injection. Colored symbols indicate mean ± SEM. Grey symbols indicate individual values.

### Participation of lipogenesis in Agrp neuron-mediated shifts in metabolism

A physiological scenario in which the metabolic shift promoted by Agrp neuron activation is expected to be important is during positive energy balance (i.e., a metabolic state coupled to weight gain), as favoring carbohydrate utilization would allow storage of lipids and re-route of energy substrates to undergo de novo lipogenesis, further increasing fat deposition. In line with this, we found that, in absence of food ingestion, Agrp neuron activation decreased circulating levels of non-esterified fatty acids (NEFAs, Figure 5B) with no changes in blood glucose levels (Figure 5C) in well-fed mice (Figure 5A). Because circulating NEFAs decrease upon Agrp neuron activation, these results suggest a decrease in release, and possibly an increase in deposition. To test this hypothesis, we measured expression levels of genes involved in lipid metabolism in the white adipose tissue (WAT) from *Agrp*^Trpv1^ and control mice 60 minutes after capsaicin injection. We found a decrease in the expression level of *Ppara* (Figure 5D), a gene involved in the promotion of fat catabolism. We also found a significant increase in expression levels of *hexokinase2* (*hk2*; Figure 5D), a rate-limiting enzyme involved in glycolysis, a critical metabolic step to provide carbons for de novo lipogenesis. Hormone-sensitive lipase (HSL) is an essential step in the breakdown of triglycerides to release fatty acids in circulation. Activation of HSP occurs by phosphorylation of this enzyme in several serine residues. Upon activation of Agrp neurons, we found decreased levels of phosphorylated HSL (pHSL) in WAT compartments (Figure 5E). Together, these experiments indicate activation of Agrp neurons leads to increased lipogenesis and decreased lipolysis in the WAT.

**Figure 5:**
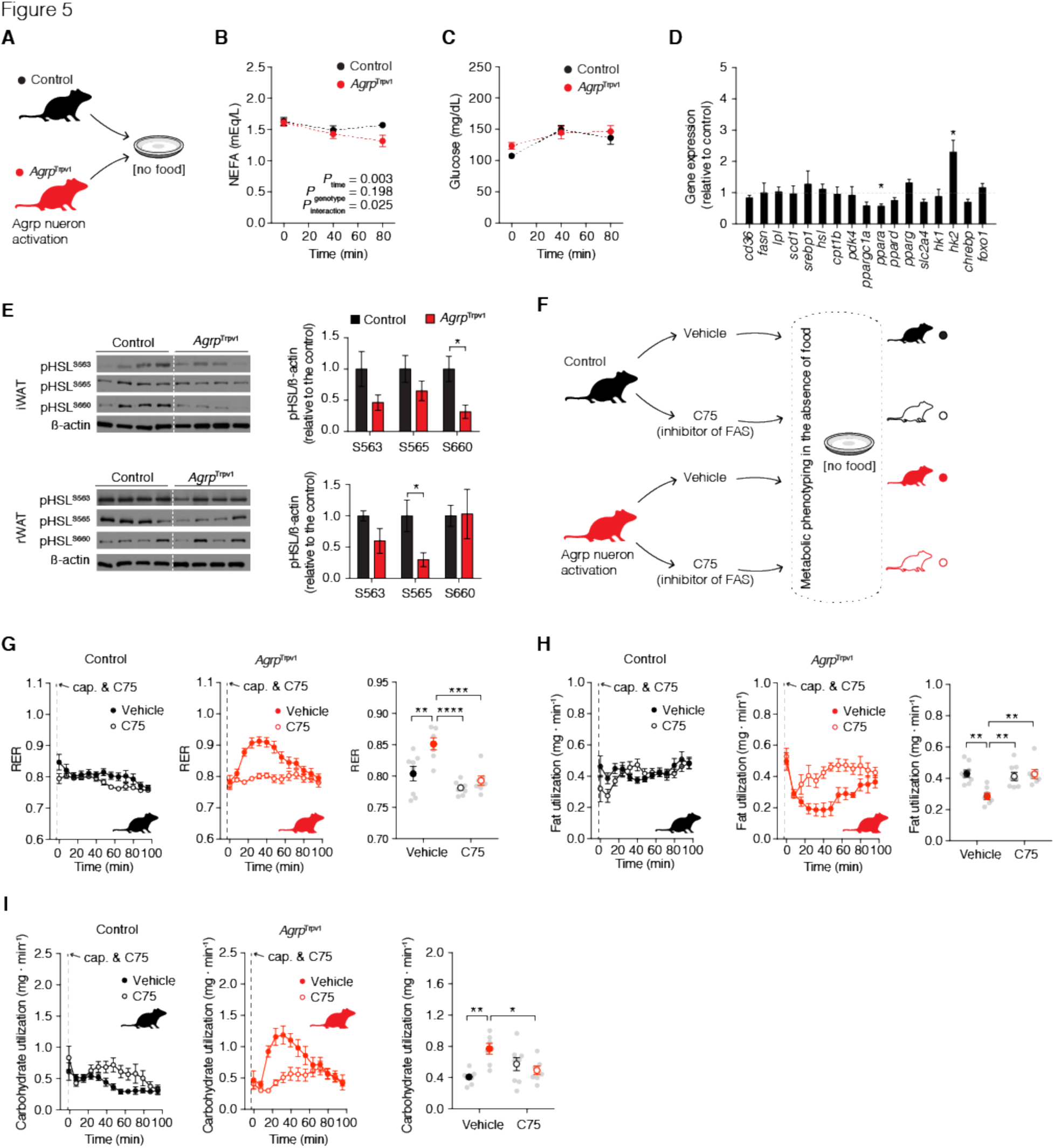
Agrp neurons promote lipogenesis. (**A**) Well-fed control (in black) and *Agrp*^Trpv1^ (in red) mice were injected with capsaicin (10 mg/kg, i.p.). One group of animals was tail bled before and after (40 and 80 min) injection to measure blood levels of NEFA and glucose. Different groups of animals were sacrificed 60 minutes after capsaicin injection and white adipose tissue (WAT) was collected for biochemical analyses. (**B**) NEFA levels (interaction: *F*_2, 54_ = 3.91, *P* = 0.02; time: *F*_2, 54_ = 9.64, *P* = 0.0003; genotype: *F*_1, 27_ = 1.74, *P* = 0.19; MCT: *P* 80 min = 0.03). (**C**) Blood glucose levels (interaction: *F*_2, 54_ = 1.47, *P* = 0.23; time: *F*_2, 54_ = 14.6, *P* < 0.0001; genotype: *F*_1, 27_ = 0.67, *P* = 0.41; MCT: not significant). In B and C: control (n = 15) and *Agrp*^Trpv1^ mice (n = 14). Statistical analysis was performed using two-way ANOVA with time as a repeated measure followed by Holm-Sidak’s multiple comparisons test (MCT). (**D**) Analysis of the transcriptional profile of the WAT upon activation of Agrp neurons. Data are normalized to control levels (dashed line). Statistical analysis was performed using student’s *t*-test. * *P* < 0.05. (**E**) Western blotting analysis of phosphorylated HSL in inguinal WAT and retroperitoneal WAT (n = 4 mice per group). Statistical analysis was performed using student’s *t*-test. * *P* < 0.05. (**F**) Experimental design to test the participation of fatty acid synthetase (FAS) in the effects of Agrp neurons on substrate utilization. Control and *Agrp*^Trpv1^ mice were randomized to receive vehicle or the FAS inhibitor (C75, 10 mg/kg, i.p.) immediately before capsaicin injection. (**G**) RER (interaction: *F*_1, 26_ = 4.36, *P* = 0.04; drug: *F*_1, 26_ = 22.28, *P* < 0.0001; genotype: *F*_1, 26_ = 11.73, *P* = 0.002). (**H**) Fat utilization (interaction: *F*_1, 26_ = 9.97, *P* = 0.04; drug: *F*_1, 26_ = 6.00, *P* = 0.02; genotype: *F*_1, 26_ = 6.56, *P* = 0.01). (**I**) Carbohydrate utilization: (interaction: *F*_1, 26_ = 12.18, *P* = 0.001; drug: *F*_1, 26_ = 0.76, *P* = 0.38; genotype: *F*_1, 26_ = 4.90, *P* = 0.03). In G-I, statistical analysis was performed using two-way ANOVA on the mean response after drug and capsaicin injection; genotype (control vs. *Agrp*^Trpv1^) and drug (vehicle vs. C75) were used as factors for the ANOVA. Holm-Sidak’s multiple comparisons test (MCT) was used to find post-hoc differences among groups. MCTs are indicated as * *P* < 0.05, ** *P* < 0.01, *** *P* < 0.001, and **** *P* < 0.0001 in figure panels. Control mice + vehicle (n = 8); control mice + C75 (n = 8); *Agrp*^Trpv1^ mice + vehicle (n = 7); *Agrp*^Trpv1^ mice + C75 (n = 7). Dashed line indicates time of injections. Colored symbols indicate mean ± SEM. Grey symbols indicate individual values.

De novo lipogenesis can drive RER above 1.0 (Frayn, 1983), and could be a potential factor involved in Agrp neuron mediated acute shifts in RER. Additionally, in our dietary clamp experiments (Figure 2), we found negative results for fat utilization in mice that ate large amounts of sugars (Figure 2 supplement 1F). Because calculated fat utilization is the sum of true rates of fat oxidation minus the rate of synthesis of fat from carbohydrates (de novo lipogenesis) (Frayn, 1983), an increase in synthesis leads to a net decrease in calculated fat utilization. Thus, negative values for calculated fat utilization *only* occur when the rate of synthesis is higher than the rate of oxidation (Frayn, 1983), and are pathognomonic of ongoing lipid synthesis. To test for the participation of fat synthesis in the rapid effects of Agrp neurons on metabolism, we blocked fatty acid synthetase (FAS), a key enzyme involved in fat storage (Lodhi et al., 2012). We treated mice with a pharmacological inhibitor of FAS (C75, 10 mg/kg, i.p.) (Kuhajda et al., 2000; Loftus et al., 2000) and activated Agrp neurons in indirect calorimetry chambers (Figure 5F). Treatment of control mice with the FAS inhibitor had no effects on RER (Figure 5G), fat utilization (Figure 5H) or carbohydrate utilization (Figure 5I). The lack of effects of FAS inhibition on substrate utilization is in line with the low levels of lipogenesis during the light cycle of mice. In contrast to control animals, inhibition of FAS blocked the effects of Agrp neuron activation on substrate utilization (Figures 5G-I). These results demonstrate Agrp neurons rapidly shift metabolism towards lipogenesis in well-fed animals.

### Sympathetic signaling mediates peripheral effects of Agrp neurons

We next determined if Agrp neurons control peripheral substrate utilization via SNS signaling. Norepinephrine release and binding to adrenergic receptors on fat compartments promotes lipolysis, while its inhibition favors lipogenesis. Accordingly, we predicted that Agrp neurons control adiposity by inhibiting sympathetic signaling on fat compartments and promoting lipogenesis in anabolic states. To test for this hypothesis, we treated mice with a ß3-adrenergic receptor agonist (CL 316,243) (Bloom et al., 1992) (Figure 6A), as ß3-adrenergic receptors are highly selective to fat compartments (Grujic et al., 1997). Treatment of control mice with CL 316,243 did not alter RER (Figure 6B), but prevented the increase in RER upon Agrp neuron activation in *Agrp*^Trpv1^ mice (Figure 6B). When we calculated fat utilization, CL 316,243 completely reverted the inhibition of whole body fat utilization upon Agrp neuron activation in *Agrp*^Trpv1^ mice but had no effects in control animals (Figure 6C). Concomitantly, CL 316,243 prevented the increase in carbohydrate utilization upon activation of Agrp neurons (Figure 6D). These results demonstrate that promotion of ß3-adrenergic receptor signaling in fat compartments is sufficient to revert the effects of Agrp neurons on peripheral fuel metabolism.

**Figure 6:**
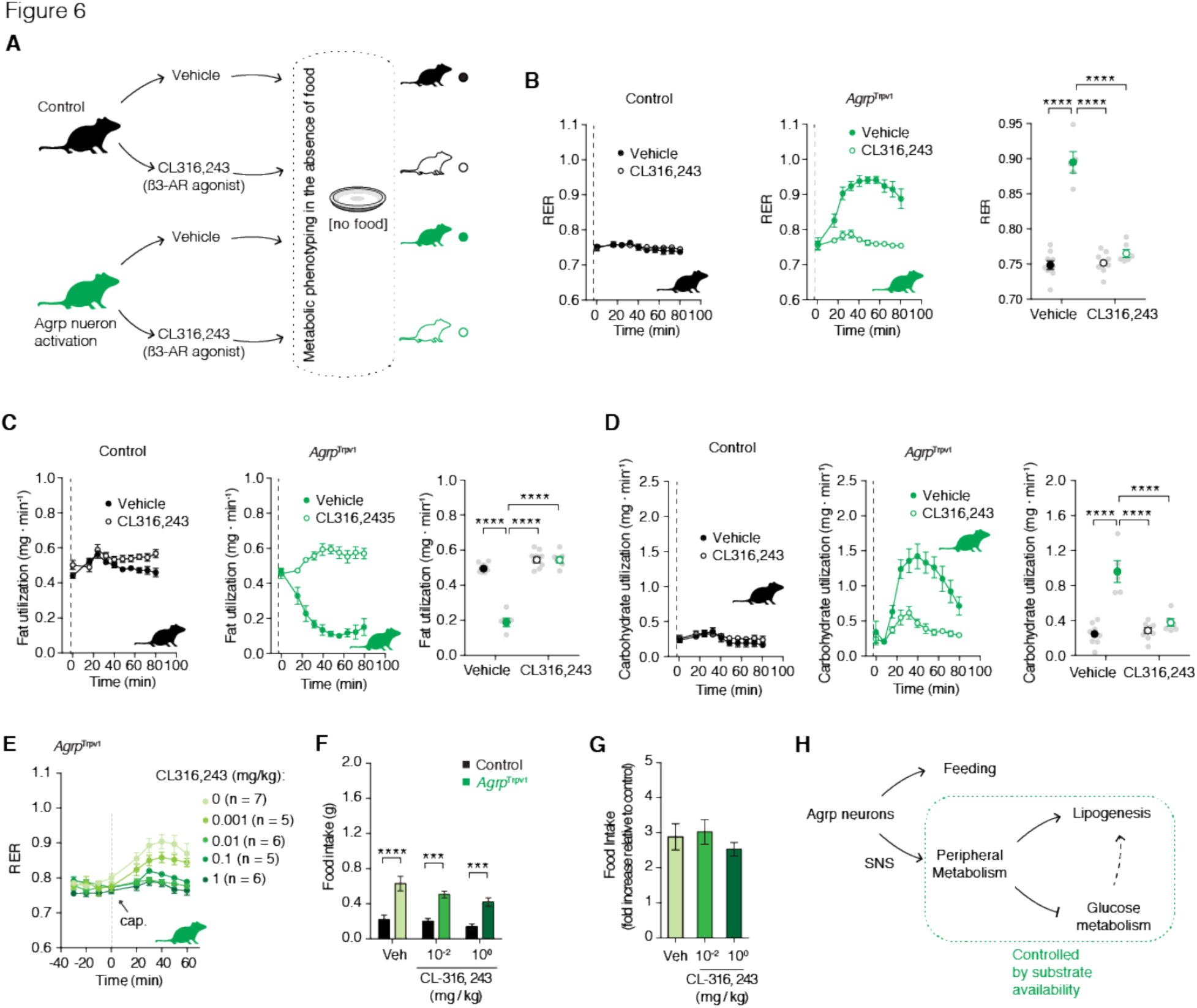
SNS signaling is involved in peripheral effects of Agrp neurons. (**A**) Experimental design to test the participation of the sympathetic nervous system (SNS) in the effects of Agrp neurons on substrate utilization. Control (black) and *Agrp*^Trpv1^ mice (green) were randomized to receive vehicle or the ß3-adrenergic receptor agonist (CL316,243, 1 mg/kg, i.p.); vehicle or CL316,243 were injected immediately before capsaicin. (**B**) RER (interaction: *F*_1, 25_ = 75.09, *P* < 0.0001; drug: *F*_1, 25_ = 67.96, *P* < 0.0001; genotype: *F*_1, 25_ = 108.4, *P* < 0.0001). (**C**) Fat utilization (interaction: *F*_1, 25_ = 85.56, *P* < 0.0001; drug: *F*_1, 25_ = 150.4, *P* < 0.0001; genotype: *F*_1, 25_ = 86.42, *P* < 0.0001). (**D**) Carbohydrate utilization: (interaction: *F*_1, 25_ = 29.74, *P* < 0.0001; drug: *F*_1, 25_ = 22.75, *P* < 0.0001; genotype: *F*_1, 25_ = 50.12, *P* < 0.0001). In B-D, statistical analysis was performed using two-way ANOVA on the mean response after drug and capsaicin injection; genotype (control vs. *Agrp*^Trpv1^) and drug (vehicle vs. CL316,243) were used as factors for the ANOVA. Holm-Sidak’s multiple comparisons test (MCT) was used to find post-hoc differences among groups. MCTs are indicated as * *P* < 0.05, ** *P* < 0.01, *** *P* < 0.001, and **** *P* < 0.0001 in figure panels. Control mice + vehicle (n = 9); control mice + CL316,243 (n = 9); *Agrp*^Trpv1^ mice + vehicle (n = 5); *Agrp*^Trpv1^ mice + CL316,243 (n = 6). Dashed line indicates time of injections. Colored symbols indicate mean ± SEM. Grey symbols indicate individual values. (**E**) Dose-response of CL316,243 injected immediately before capsaicin in *Agrp*^Trpv1^ mice (dashed line denotes injection time). Number of animals used per experimental group shown in the panel. (**F**) Food intake response of control and *Agrp*^Trpv1^ mice to activation of Agrp neurons when injected with different doses of CL316,243 (n = 7 for all groups); interaction (*F*_2, 36_ = 0.95, *P* = 0.39); drug (*F*_2, 36_ = 4.16, *P* = 0.02); genotype (*F*_1, 36_ = 64.96, *P* < 0.0001). Statistical analysis was performed using two-way ANOVA with genotype and drug as factors. Holm-Sidak’s multiple comparisons test (MCT) was used to find post-hoc differences genotypes. MCTs are indicated as *** *P* < 0.001 and **** *P* < 0.0001 in figure panel. (**G**) Related to F, but fold change in food intake in *Agrp*^Trpv1^ related to control mice. Bars and symbols indicate mean ± SEM. (**H**) Diagram illustrating the model for different roles of Agrp neurons in the control of feeding and peripheral metabolism. Agrp neurons control peripheral metabolism via the SNS, but switching substrate utilization towards use of carbohydrate.

Because Agrp neurons are characteristically linked to food intake control (Aponte et al., 2011; Gropp et al., 2005; Hahn et al., 1998; Krashes et al., 2011; Luquet et al., 2005; Rossi et al., 1998), we investigated whether the effects of Agrp neurons on fuel utilization are mechanistically linked to feeding. Thus, we tested whether CL 316,243 could acutely block the effects of Agrp neuron activation on food intake. We first performed a dose response study to detect the range of CL 316,243 doses that could revert the increase in RER upon Agrp neuron activation. We found that doses as low as 0.01 mg/kg almost completely reverted the effects of Agrp neurons on RER (Figure 6E). Next, we selected two doses of CL 316,243 (1.00 and 0.01 mg/kg) to investigate its effects on Agrp neuron mediated food intake. CL 316,243 is highly anorexigenic (Grujic et al., 1997), but its effects are observed after several hours and not as rapid as the effects of Agrp neurons on feeding (Aponte et al., 2011; Dietrich et al., 2015; Krashes et al., 2011). In all conditions tested, CL 316,243 did not revert the effects of Agrp neuron activation on food intake (Figures 6F-G). These latter findings indicate a divergence between the feeding and metabolic mechanisms underlying Agrp neuron function and support the argument that Agrp neurons favor the storage of fat (lipogenesis) in situations of energy surfeit (Figure 6H).

## Discussion

We reported here a mechanism by which elevated Agrp neuron activity shifts metabolism towards lipid storage via the SNS. The SNS releases norepinephrine in target organs to regulate a variety of physiological functions. In WAT, which stores excess of energy in the form of fat, activation of the SNS leads to lipolysis (Bartness et al., 2014; Correll, 1963; Zeng et al., 2015), breaking down triglycerides to release free fatty acids in the circulation (Zeng et al., 2015). Lipolysis is critical for survival during periods of food scarcity, with fatty acids becoming the predominant energy substrate. Conversely, inhibition of lipolysis favors lipogenesis, leading to fat deposition (Rutkowski et al., 2015). This is an important adaptive response to allow storage of the excess of energy for later mobilization during food deprivation. However, when this anabolic state (lipogenesis > lipolysis) is sustained it can become maladaptive and trigger obesity. Here, we provided evidence that elevated activity of Agrp neurons is sufficient to promote this shift in metabolism towards storage of nutrients (largely carbohydrates) as fat.

Our results are in apparent contrast with a previous publication showing that neonatal ablation of Agrp neurons leads to decreased RER, increased metabolic efficiency and obesity in adult animals (Joly-Amado et al., 2012). Contrary to adult ablation of Agrp neurons which leads to cessation of feeding and death (Gropp et al., 2005; Luquet et al., 2005; Wu and Palmiter, 2011), neonatal ablation of these neurons is compatible with life (Luquet et al., 2005). In the study by (Joly-Amado et al., 2012), Agrp neurons were ablated during the first postnatal week. The authors found Agrp neuron-ablated mice presented metabolic abnormalities only later in adulthood (> 3 months of age), which were in similar direction to acute activation of Agrp neurons in adult animals (this report and (Srisai, 2016; Steculorum et al., 2016)). These results strengthen the idea that Agrp neurons have distinct roles at different developmental stages (Dietrich et al., 2012), and that altering the function of Agrp neurons early in life leads to compensatory mechanisms with long lasting physiological implications.

Recent studies have described the *in vivo* dynamics of Agrp neuron activity (Betley et al., 2015; Chen et al., 2015; Mandelblat-Cerf et al., 2015). However, not all cells were homogeneous in their response. While 2/3 of Agrp neurons had decreased activity upon presentation of food/eating, the other third did not change firing rate or even increased activity during eating. The fact that Agrp neurons had heterogeneous response to feeding further highlights the existence of subpopulations of Agrp neurons that are functionally, anatomically (Betley et al., 2013; Padilla et al., 2016; Steculorum et al., 2016), and potentially genetically distinct. In light of our data, it is thus possible that the sustained activity of distinct subpopulation of Agrp neurons during feeding/refeeding might operate to maximize energy storage by shifting substrate utilization towards lipogenesis.

Less expected than the elevated activity of Agrp neurons during food deprivation were the findings that during diet-induced obesity (DIO) the activity of these neurons is also elevated (Baver et al., 2014; Diano et al., 2011; Dietrich et al., 2013; Wei et al., 2015). In a previous study, we have identified that the elevated activity of Agrp neurons during high-fat feeding relies on intracellular mitochondria fusion machinery (Dietrich et al., 2013). We found that deletion of mitofusins selectively in Agrp neurons was sufficient to blunt the increase in neuronal activity in response to a high-fat diet (HFD); as a consequence, mice were resistant to DIO. Intriguingly, we found only minor or no effects of these genetic manipulations on food intake (Dietrich et al., 2013), indicating that, in obesogenic conditions, Agrp neurons also control other aspects of physiology that are independent of food consumption. The elevated activity of Agrp neurons during obesity development (Baver et al., 2014; Diano et al., 2011; Dietrich et al., 2013; Wei et al., 2015) could be involved in the metabolic shifts towards fat deposition (lipogenesis) as observed here. This consideration, however, will require further testing in future studies.

In summary, we showed that Agrp neurons in the hypothalamus rapidly control whole-body nutrient utilization by shifting metabolism towards lipogenesis via the sympathetic nervous system. These studies expanded the repertoire of functions attributed to Agrp neurons, which emerge as controllers of a variety of physiological functions in addition to food intake. Because there are several thousands of Agrp neurons in the mammalian brain, future studies will be important to determine whether each subpopulation of Agrp neurons regulate a specific physiological function. Because Agrp neurons release several molecules, including AGRP, NPY, and GABA, it is also possible that this multifaceted function of Agrp neurons arises from the combination of released molecules. These findings have strong implications for our understanding of how neuronal circuits involved in metabolism regulation function and, consequently, to our understanding of severe disordered conditions such as obesity.

## Experimental Procedures

### Animals

Mice used in the experiments were 3–8 months old from both genders. *Agrp*^Trpv1^ mice were: *Agrp*^Cre/+^:: *Trpv1*^KO/KO^:: *R26-LSL-Trpv1*^Gt/+^; control animals were either *Agrp*^Trpv1^ mice injected with vehicle (3.3% Tween 80 in saline) or *Trpv1*^KO/KO^: *R26-LSL-Trpv1*^Gt/+^ mice injected with capsaicin. All animals were littermates (Agrp neuron activated and controls) in the experiments. We did not observe any differences between the two control groups and, therefore, throughout the manuscript we referred to them as “controls”. We have carefully characterized this animal model to activate Agrp neurons and reported elsewhere (Dietrich et al., 2015; Ruan et al., 2014). We have performed dose-response curves for capsaicin and identified the dose of 10 mg/kg (i.p.) as optimal to induce behavior phenotypes in *Agrp*^Trpv1^ mice. We have also performed a dose-response of capsaicin and measured changes in RER (1, 3, 10 and 30 mg/kg, i.p.; experiments not reported). We also found that 10 mg/kg was the optimal dose to promote changes in RER. Thus, we selected this dose of capsaicin for our studies.

The following mouse lines were used in this study: Agrptm1(cre)Lowl/J, Gt(ROSA)26Sortm1(Trpv1,ECFP)Mde/J, Trpv1tm1Jul/J. These lines are available from The Jackson laboratories. All animals were kept in temperature and humidity controlled rooms, in a 12/12h light/dark cycle, with lights on from 7:00AM-7:00PM. Food and water were provided ad libitum, unless otherwise stated. All procedures were approved by IACUC (Yale University).

### Drugs

The following compounds were used in the reported studies: C75 (RPMI medium 1640; from Tocris); capsaicin (3.3% Tween-80 in saline; from Sigma), and CL-316, 245 (in PBS; from Tocris). All drugs were injected in a volume of 10 ml/kg of body weight intraperitoneally (i.p.). When multiple injections were performed in the same experiment, the volume of each injection was adjusted to a total volume of 10 ml/kg per animal.

### Metabolic Assays and Biochemical Analysis

For all experiments animals were housed in individual cages at least three days prior to the experiment. Blood samples were collected from the tail in order to measure glucose and free fatty acids (NEFA) levels. Glucose was measured using a One Touch Ultra 2 glucometer. After blood centrifugation, serum was collected and used to measure NEFA as indicated by manufacturer (WAKO, Japan).

### Gene Expression and Western blotting

Animals were deeply anesthetized with ketamine and xylazine and euthanized by decapitation. Tissues were collected and frozen in liquid nitrogen. Tissues were lysed in buffer containing 1% Nonidet P-40, 50 mM Tris 3 HCl, 0.1 mM EDTA, 150 mM NaCl, proteinase inhibitors and protein phosphatase inhibitors. Equal amounts of protein lysate were electrophoresed on SDS-PAGE gels and transferred to PVDF membranes. Primary antibodies (Lipolysis Activation Antibody Sampler Kit #8334, Cell Signaling) were incubated at 4°C overnight. Membranes were washed and incubated with secondary antibodies conjugated to horseradish peroxidase. Protein levels were visualized using ECL chemiluminescent substrate and quantified using ImageJ.

Total RNA was extracted from mouse tissues using RNeasy^®^ lipid mini kit (Qiagen). cDNA was reverse transcribed (Bio-Rad) and amplified with SYBR Green Supermix (Bio-Rad) using a Light Cycler 480 real-time PCR system (Roche). Data were normalized to the expression of *Actin*, *Gusb* and *Arbp*. Primer sequences are available on request.

### Indirect calorimetry

Oxygen consumption (VO_2_) and CO_2_ production (VCO_2_) were measured in four to eight mice simultaneously in indirect calorimetry chambers (TSE Systems, Germany). Measurements were recorded every 8-12 minutes over the entire course of the experiment (except for the experiment in which ad libitum food intake was measured during one entire day). Respiratory exchange ratio (RER) was calculated as the ratio between VCO_2_ and VO_2_. Whole body fat utilization was calculated using the follow equation: 1.67 * (VO_2_-VCO_2_). Whole body carbohydrate utilization was calculate using the follow equation: 4.55 * VCO_2_ – 3.21 * VO_2_ (Frayn, 1983). All animals were single housed during the experiments in calorimetry chambers. For the experiments in which different diets were fed to the animals, the following diets were used: low-fat high-sugar diet (LFHS; D12450B, Open Source Diets, USA); high-fat diet 45% calories from fat (HF45; D12451, Open Source Diets, USA); and high-fat diet 60% calories from fat (HF60; D12492, Open Source Diets, USA). For the glucose response study, mice were provided with glucose (D-Glucose, G8270, Sigma, USA) via gavage. We used saline as vehicle and three doses of glucose (1, 2 and 3 g/kg body weight, via gavage feeding). Food was removed 2 hours before the experiment during the light cycle of the animals. Calorimetry was recorded during 60 minutes prior injection of capsaicin and diet switch. For the experiments that no food was provided, food was removed from the cages 2 hours before injecting mice with capsaicin/vehicle. Baseline calorimetry was recorded for 60 minutes, and then the effects of Agrp neuron activation were recorded for 60-120 minutes. Similar procedures were used in the experiments in which compounds were given to mice prior the experiment. Glucose (2 g/kg, via gavage), C75 (10 mg/kg, i.p.) or CL-316,243 (0.01-1.00 mg/kg, i.p) were given together with capsaicin. In all cases, the drugs were injected immediately before capsaicin using two different syringes. Total injected volume was adjusted for the maximal dose of 10 ml/kg mouse body weight.

### Statistical Considerations

Matlab R2016a, PASW Statistics 18.0, Prism 7.0 and Adobe Illustrator CS6/CC were used to analyze data and plot figures. Student’s *t* test was used to compare two groups. ANOVA was used to compare multiple groups. When necessary, multiple comparisons post hoc test (MCT) was used (Holm-Sidak’s test). When homogeneity was not assumed, the Kruskal–Wallis nonparametric ANOVA was selected for multiple statistical comparisons. The Mann–Whitney U test was used to determine significance between groups. Statistical data are provided in the figures. *P* < 0.05 was considered statistically significant.

## Acknowledgments

We thank Matthew Rodeheffer, Ivan de Araujo, Hai-bin Ruan and Xiaoyong Yang for comments on this project and manuscript. M.O.D. received support from Brain and Behavior Research Foundation (NARSAD Young Investigator Award), Yale Center for Clinical Investigation Scholars Award (NCATS, UL1 TR000142), National Institute Of Diabetes And Digestive And Kidney Diseases (1R01DK107916-01), DRC (P30 DK045735), Whitehall Foundation, Charles H. Hood Foundation, CNPq (487096/2013-4 and 401476/2012-0, Brazil) and CAPES (88881.068059/2014-01, Brazil). JPA and MRZ were partially supported by a fellowship from the Science Without Borders program (Brazil).

## Author Contributions

JB and MZ helped to perform the experiments and analyze the data. JAP and MOD performed the experiments, designed, analyzed data and wrote the manuscript.

**Figure 2 supplement 1:**
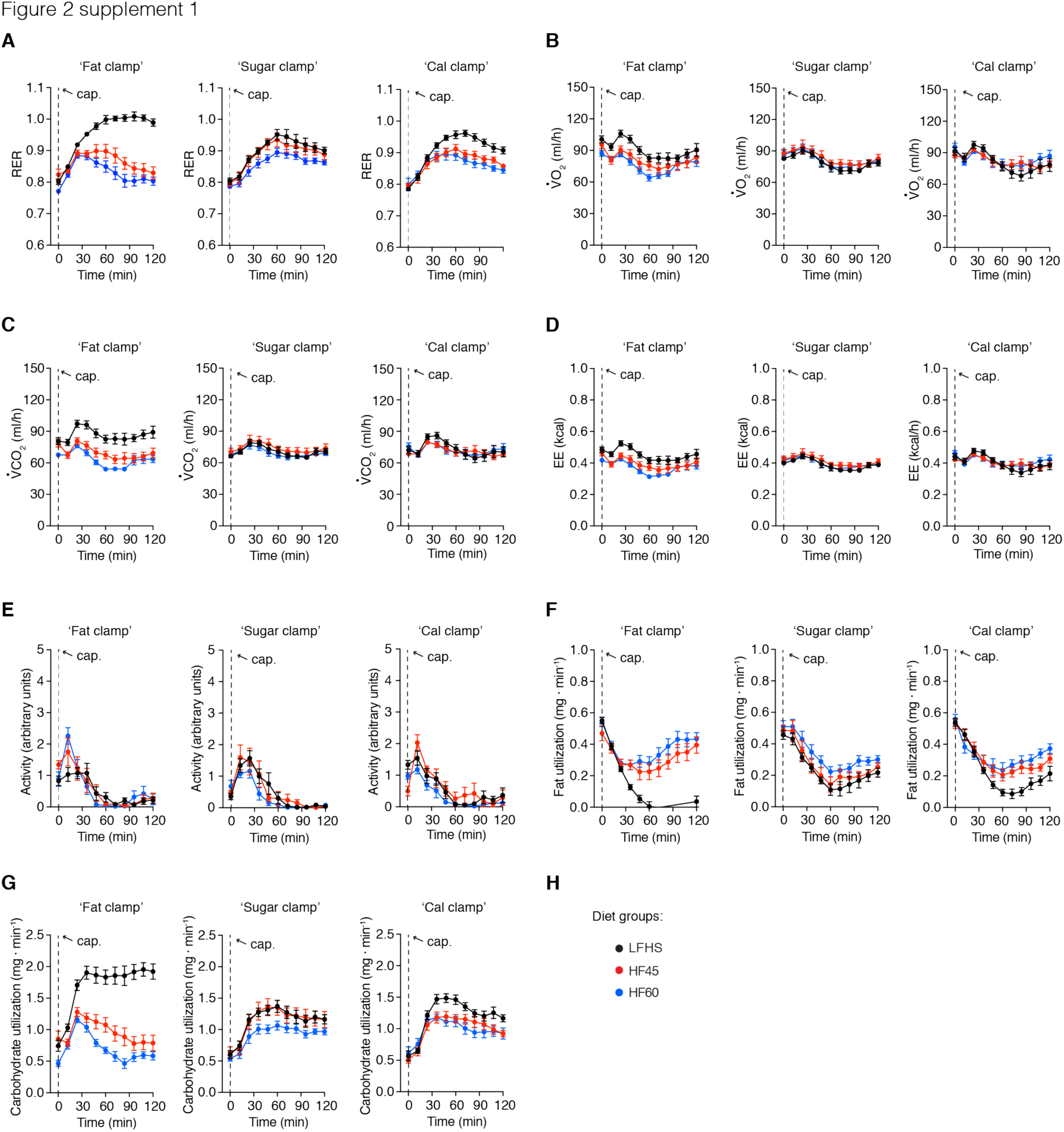
Substrate utilization in response to different diets. Detailed metabolic phenotyping data of *Agrp*^Trpv1^ mice fed different diets (Figure 2A) upon activation of Agrp neurons by peripheral injection of capsaicin (10 mg/kg, i.p.). In black, mice fed a low-fat high-sugar diet (LFHS; Research Diets D12450B); in red, mice fed a high-fat diet (45 kcal% from fat, HF45; Research Diets D12451); and in blue, mice fed a high-fat diet (60 kcal% from fat, HF60; Research Diets D12492). (**A**) RER: ‘Fat clamp’ (interaction: *F*_20, 270_ = 25, *P* < 0.0001; time: *F*_10, 270_ = 56.34, *P* < 0.0001; diets: *F*_2, 27_ = 29.13, *P* < 0.0001); ‘Sugar clamp’ (interaction: *F*_20, 270_ = 0.64, *P* = 0.87; time: *F*_10, 270_ = 65.72, *P* < 0.0001; diets: *F*_2, 27_ = 2.87, *P* = 0.07); ‘Cal clamp’ (interaction: *F*_20, 270_ = 3.51, *P* < 0.0001; time: *F*_10, 270_ = 49.89, *P* < 0.0001; diets: *F*_2, 27_ = 8.92, *P* = 0.001). (**B**) VO_2_: ‘Fat clamp’ (interaction: *F*_20, 270_ = 0.91, *P* = 0.56; time: *F*_10, 270_ = 16.08, *P* < 0.0001; diets: *F*_2, 27_ = 6.28, *P* < 0.0001); ‘Sugar clamp’ (interaction: *F*_20, 270_ = 0.18, *P* > 0.99; time: *F*_10, 270_ = 14.66, *P* < 0.0001; diets: *F*_2, 27_ = 0.69, *P* = 0.50); ‘Cal clamp’ (interaction: *F*_20, 270_ = 1.65, *P* = 0.04; time: *F*_10, 270_ = 13.98, *P* < 0.0001; diets: *F*_2, 27_ = 0.13, *P* = 0.87). (**C**) VCO_2_: ‘Fat clamp’ (interaction: *F*_20, 270_ = 1.70, *P* = 0.03; time: *F*_10, 270_ = 11.55, *P* < 0.0001; diets: *F*_2, 27_ = 22.47, *P* < 0.0001); ‘Sugar clamp’ (interaction: *F*_20, 270_ = 0.28, *P* = 0.99; time: *F*_10, 270_ = 8.59, *P* < 0.0001; diets: *F*_2, 27_ = 0.90, *P* = 0.41); ‘Cal clamp’ (interaction: *F*_20, 270_ = 1.30, *P* = 0.17; time: *F*_10, 270_ = 10.33, *P* < 0.0001; diets: *F*_2, 27_ = 0.14, *P* = 0.86). (**D**) Energy expenditure: ‘Fat clamp’ (interaction: *F*_20, 270_ = 0.84, *P* = 0.66; time: *F*_10, 270_ = 14.62, *P* < 0.0001; diets: *F*_2, 27_ = 8.92, *P* = 0.001); ‘Sugar clamp’ (interaction: *F*_20, 270_ = 0.19, *P* > 0.99; time: *F*_10, 270_ = 12.9, *P* < 0.0001; diets: *F*_2, 27_ = 0.67, *P* = 0.51); ‘Cal clamp’ (interaction: *F*_20, 270_ = 1.55, *P* = 0.06; time: *F*_10, 270_ = 12.66, *P* < 0.0001; diets: *F*_2, 27_ = 0.08, *P* = 0.91). (**E**) Ambulatory activity: ‘Fat clamp’ (interaction: *F*_20, 270_ = 1.69, *P* = 0.03; time: *F*_10, 270_ = 25.32, *P* < 0.0001; diets: *F*_2, 27_ = 0.23, *P* = 0.78); ‘Sugar clamp’ (interaction: *F*_20, 270_ = 1.17, *P* = 0.27; time: *F*_10, 270_ = 31.81, *P* < 0.0001; diets: *F*_2, 27_ = 1.01, *P* = 0.37); ‘Cal clamp’ (interaction: *F*_20, 270_ = 1.68, *P* = 0.03; time: *F*_10, 270_ = 26.55, *P* < 0.0001; diets: *F*_2, 27_ = 3.03, *P* = 0.06). (**F**) Fat utilization: ‘Fat clamp’ (interaction: *F*_20, 270_ = 19.24, *P* < 0.0001; time: *F*_10, 270_ = 63.42, *P* < 0.0001; diets: *F*_2, 27_ = 16.5, *P* < 0.0001); ‘Sugar clamp’ (interaction: *F*_20, 270_ = 0.34, *P* = 0.99; time: *F*_10, 270_ = 55.74, *P* < 0.0001; diets: *F*_2, 27_ = 2.65, *P* = 0.08); ‘Cal clamp’ (interaction: *F*_20, 270_ = 3.22, *P* < 0.0001; time: *F*_10, 270_ = 55.86, *P* < 0.0001; diets: *F*_2, 27_ = 4.76, *P* = 0.01). (**G**) Carbohydrate utilization: ‘Fat clamp’ (interaction: *F*_20, 270_ = 14.97, *P* < 0.0001; time: *F*_10, 270_ = 30.32, *P* < 0.0001; diets: *F*_2, 27_ = 45.82, *P* < 0.0001); ‘Sugar clamp’ (interaction: *F*_20, 270_ = 0.67, *P* = 0.84; time: *F*_10, 270_ = 44.44, *P* < 0.0001; diets: *F*_2, 27_ = 2.62, *P* = 0.09); ‘Cal clamp’ (interaction: *F*_20, 270_ = 2.13, *P* = 0.003; time: *F*_10, 270_ = 44.43, *P* < 0.0001; diets: *F*_2, 27_ = 5.52, *P* = 0.009). Statistical analysis was performed using two-way ANOVA with time as a repeated measure followed by Holm-Sidak’s multiple comparisons test (MCT). MCTs are not shown. Dashed line indicates time of capsaicin injection. A total n = 10 mice were used for each condition. Symbols indicate mean ± SEM.

**Figure 4 supplement 1:**
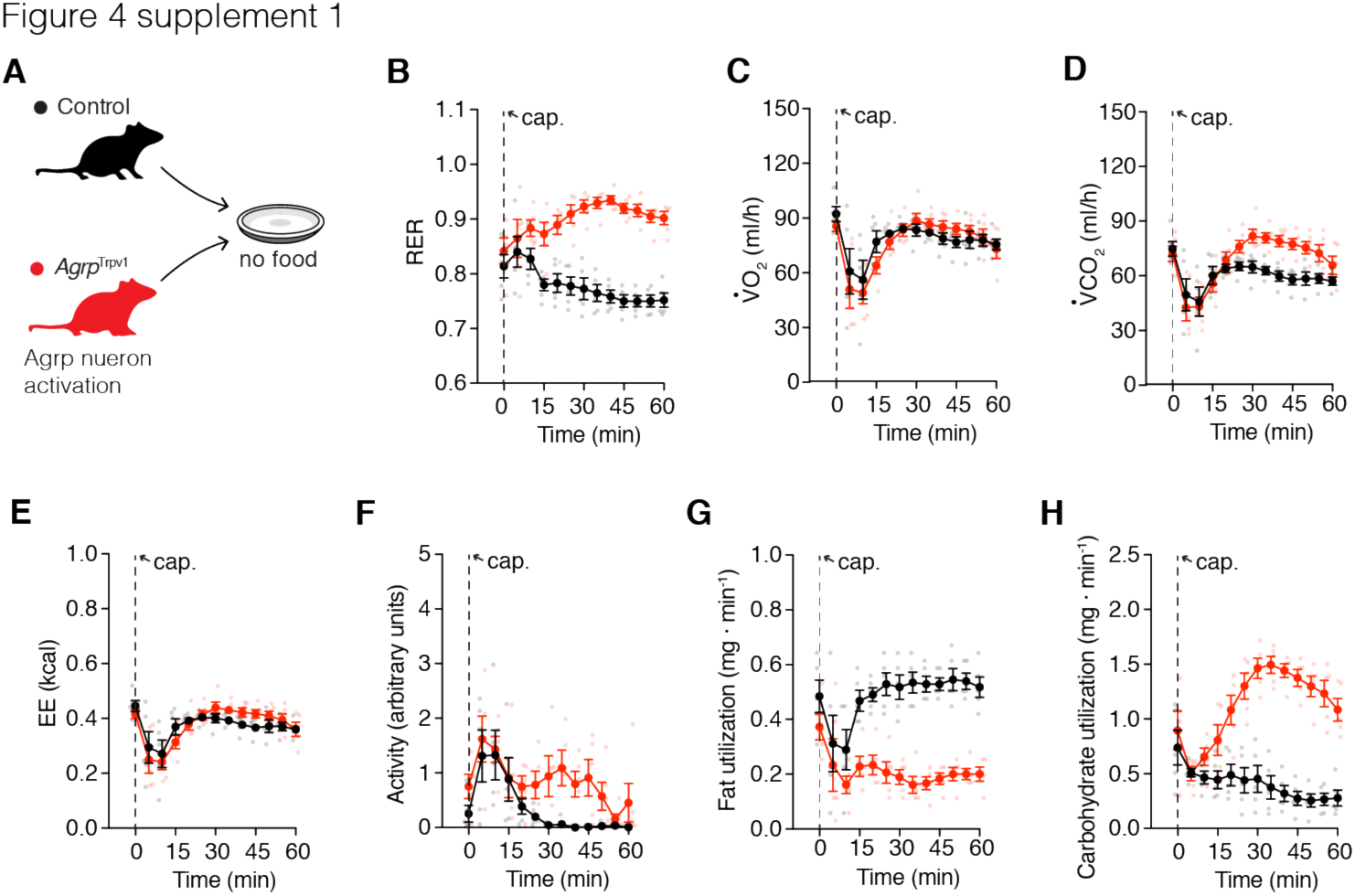
Agrp neurons control substrate utilization independently of ingestion. (**A**) Well-fed control (in black) and *Agrp*^Trpv1^ (in red) mice were tested in metabolic chambers upon injection of capsaicin without food provided. (**B**) RER (interaction: *F*_12, 96_ = 8.96, *P* < 0.0001; time: *F*_12, 96_ = 1.31, *P* = 0.22; genotype: *F*_1, 8_ = 54.45, *P* < 0.0001). (**C**) VO_2_ (interaction: *F*_12, 96_ = 0.87, *P* = 0.57; time: *F*_12, 96_ = 9.63, *P* < 0.0001; genotype: *F*_1, 8_ = 0.15, *P* = 0.70). (**D**) VCO_2_ (interaction: *F*_12, 96_ = 2.89, *P* = 0.001; time: *F*_12, 96_ = 10.66, *P* < 0.0001; genotype: *F*_1, 8_ = 10.39, *P* = 0.01). (**E**) Energy expenditure (interaction: *F*_12, 96_ = 1.16, *P* = 0.31; time: *F*_12, 96_ = 9.89, *P* < 0.0001; genotype: *F*_1, 8_ = 0.13, *P* = 0.72). (**F**) Ambulatory activity (interaction: *F*_12, 96_ = 1.12, *P* = 0.34; time: *F*_12, 96_ = 7.13, *P* < 0.0001; genotype: *F*_1, 8_ = 5.29, *P* = 0.05). (**G**) Calculated fat utilization (interaction: *F*_12, 96_ = 3.52, *P* = 0.0002; time: *F*_12, 96_ = 3.23, *P* = 0.0006; genotype: *F*_1, 8_ = 55.31, *P* < 0.0001). (**H**) Calculated carbohydrate utilization (interaction: *F*_12, 96_ = 13.52, *P* < 0.0001; time: *F*_12, 96_ = 6.25, *P* < 0.0001; genotype: *F*_1, 8_ = 58.16, *P* < 0.0001). Statistical analysis was performed using two-way ANOVA with time as a repeated measure followed by Holm-Sidak’s multiple comparisons test (MCT). MCTs are not shown. Dashed line indicates time of capsaicin injection. Small symbols indicate individual values. Large symbols indicate mean ± SEM.

